# Droplet-based microfluidics as a media optimization tool for cyanobacteria

**DOI:** 10.1101/2022.07.03.498595

**Authors:** Jialan Cao, David A. Russo, Ting Xie, G. Alexander Groß, Julie A. Z. Zedler

## Abstract

The urgent need to increase sustainability in biotechnology has led to an increased interest in photosynthetic production platforms. Cyanobacteria are particularly attractive for their fast photoautotrophic growth and genetic accessibility. However, the lack of systematic strain optimization is holding back progress towards industrialization. To overcome this bottleneck, here we introduce a droplet-based microfluidics platform capable of one- (1D) and two-dimension (2D) screening of key parameters in cyanobacterial cultivation. We successfully grew three different unicellular, biotechnologically relevant cyanobacteria: *Synechocystis* sp. PCC 6803, *Synechococcus elongatus* UTEX 2973 and *Synechococcus* sp. UTEX 3154. Highly-resolved 1D screening of nitrate, phosphate, carbonate, and salt content show that nitrate and/or phosphate can be limiting in standard cultivation media. Finally, we show that 2D screening results from our microfluidic setup translate well to a laboratory scale high-density cultivation setup. This study demonstrates that droplet-based microfluidics by micro segmented-flow are a powerful tool for high-throughput optimization of cyanobacterial cultivation.

## Introduction

Cyanobacteria have attracted interest for sustainable biotechnology due to their fast photoautotrophic growth, minimal nutritional requirements, and metabolic capacity for heterologous product formation. Transfer of existing systems to an industrial scale is however still rare and yet to come of age. Despite this, the market of cyanobacterial products is rapidly growing and cyanobacteria already account for half of the European algae production^1^. The existing products, however, are largely limited to “Spirulina” (*Arthrospira* spp.) which can be grown in high pH and high salt conditions. To efficiently exploit cyanobacterial diversity, and increase their biotechnological relevance, several challenges need to be solved. One key aspect is strain-specific optimization of growth conditions. To achieve this, high-throughput screening methods that are suitable for studying cyanobacterial chassis are needed. Standard high throughput methods for heterotrophic systems, such as micro-well cultivation and live-cell monitoring systems, are often not compatible with cyanobacteria due to the requirement for controlled light exposure, extended cultivation periods and autofluorescence interference with optical measurements. In addition, increasing of throughput by reducing volume raises new challenges in terms of oxygen supply, evaporation control and homogeneity. Thus, it is of great importance to establish alternative, cost-efficient technologies that are tailored for high-throughput screening and optimization of cyanobacteria.

Microfluidics is an established tool in chemical analytics and micro reaction technology and has played a pivotal role in the development of high-throughput analysis strategies for biological systems. It was first recognized as a promising tool for biological applications more than two decades ago. Starting with micro capillary electrophoresis^2^ and micro flow-through PCR^3,4^, miniaturized fluidic techniques became of interest for fast and efficient bioanalytical procedures. The ability to miniaturize volumes and multiplex sample handling and analysis then also attracted interest in molecular biology as well as in microbiology. For cultivation, droplet-based microfluidics by micro segmented-flow was first introduced to separate soil microorganisms from complex soil microbial communities^5^. In small droplets, the principle of stochastic confinement applies. In other words, rare individuals and their released molecules can accumulate to high densities, thus, enabling single cell experiments^6,7^. The segmented-flow principle enables an easy screening of sample series with concentration gradients and allows the measurement of organismic responses to stress factors like antibiotics, nanoparticles or heavy metals^8–10^ with high resolution. In addition, this approach allows the screening of two and three-dimensional concentration spaces with a very small amount of material in a single experimental run. Thus, opening the possibility of conveniently detecting combinatorial effects of, for example, toxins, pharmaceutical drugs and nutrients^11^. Furthermore, droplet-based microfluidics is in use for detection of antibiotic resistances and for the development of new antibiotics^12^. Besides bacteria, the micro segmented-flow technique has also proven to be suitable for eukaryotic microorganisms such as *Chlorella*^13^ or even embryos of multicellular organisms^14^ and for the characterization of the concentrationdependent response of complex unknown environmental bacterial communities on toxic substances^15,16^. As applied to cyanobacterial systems, microfluidics is an emerging technology with only a handful of applications. For instance, a microfluidics-based tool for harvesting cyanobacterial biomass was developed for *Synechocystis* sp. PCC 6803^17^ (hereafter PCC 6803). In addition, microfluidics have been used for lactate productivity screening in a PCC 6803 CRISPRi library^18^ as well as to screen for ethanol production^19^.

Here, we established a droplet-based microfluidic setup that allows for high-throughput media composition optimization for biotechnologically relevant cyanobacteria. We tested the compatibility of our system with three different cyanobacterial strains using biomass accumulation as the screening parameter. Our microfluidic setup allowed for continuous cultivation and analysis of the cells where individual medium parameters such as macronutrient content as well as salt and bicarbonate concentrations were varied in small steps allowing for highly parallel, high-throughput screening. In addition, we demonstrated that our platform can screen two parameters simultaneously to explore the two-dimensional effect space of combined variables. In conclusion, we demonstrate that microfluidics can be used to improve medium composition and cultivation conditions in a cost- and time-effective manner to unlock the exploitation of biotechnologically relevant cyanobacteria.

## Materials and Methods

### Strains and Growth Conditions

Three cyanobacterial wild-type strains were used: a non-motile, glucose-tolerant substrain of *Synechocystis* sp. PCC 6803 (originally obtained from Patrik Jones, Imperial College London), *Synechococcus elongatus* UTEX 2973 (originally obtained from Himadri Pakrasi, University of Washington) (hereafter UTEX 2973) and *Synechococcus* sp. UTEX 3154 (a spontaneous mutant of *Synechococcus* sp. PCC 11901^20^ which does not require vitamin B12 for growth, obtained from the UTEX culture collection) (hereafter UTEX 3154). PCC 6803 and PCC 2973 were grown in BG-11 medium^21^ supplemented with 10 mM N-Tris(hydroxymethyl)methyl-2-aminoethanesulphonic acid (TES) buffer (pH 8.0) (BG-11 TES medium). Medium A^22^ with D7 micronutrients^23^ (AD7 medium) (pH 7.5) was used for cultivation of UTEX 3154. All cultures were maintained at 30°C with 20 – 50 μmol photons m^-2^ s^-1^ white light (Lumilux cool white L 15W/840 fluorescent lamps (Osram, Germany)) on BG-11 or AD7 medium plates supplemented with 1.5% bacto-agar. Liquid pre-cultures for inoculation of the microfluidic coils were grown in glass tubes bubbled with 3% CO_2_-supplemented air at 30°C with approximately 60 μmol photons m^-2^ s^-1^ white light for 3 to 4 days (culture volume: 20 mL).

To test improved nitrogen:phosphorus (N:P) ratios based on 2D screening experiments, standard BG-11 medium and BG-11 medium with adjusted NaNO_3_ and K_2_HPO_4_ concentrations were prepared. Liquid cultures were inoculated for PCC 6803 and UTEX 2973 in a high-density cultivation setup (HDC 6.10 starter kit CellDEG, Germany)^24^ using 25 mL culture vessels with a culture volume of 10 mL. The cultures were supplemented through a membrane with CO_2_ at a partial pressure of approximately 32 mbar (reference T = 20°C) by a carbonate buffer (3 M KHCO_3_ and 3 M K_2_CO_3_, ratio 4:1)^24^. The growth setup was placed on a Unimax 1010 orbital shaker (Heidolph Instruments, Germany) and incubated with shaking at 280 rpm at 30°C with 50 μmol photons m^-2^ s^-1^ white light (Lumilux cool white L 15W/840 fluorescent lamps (Osram, Germany)). Main cultures for the growth experiment were inoculated in triplicates from a pre-culture grown in the same medium and conditions to a starting optical density (OD) at 750 nm of 0.3 and monitored for 7 days by measuring OD at 750 nm every 24 hours using a GENESYS 10S UV-Vis Spectrophotometer (Thermo Scientific, Germany). For PCC 6803, the optimized BG-11 medium contained 0.4 mM K_2_HPO_4_ and 27 mM NaNO_3_ and for UTEX 2973 0.45 mM K_2_HPO_4_ and 30 mM NaNO_3_. All other elements of the medium were kept the same as in the standard BG-11 TES medium.

### Microfluidic Cultivation

#### Microfluidic arrangement

Details on the fluidic devices, the optical detection unit and the applied methods for realizing concentration-graded droplet sequences were reported earlier9. Here, a similar experimental setup (Fig. 1) was used for one- and two-dimensional screening of medium parameters. Briefly, the system is based on a syringe pump with six dosing units (Cetoni GmbH, Germany). The microfluidic droplets were generated by a self-developed droplet generator comprising of a 6-port manifold25. Droplets with varying composition are generated by controlled dosing of effectors, culture medium and cell suspension into a flow of carrier liquid (perfluoromethyldecalin (PP9)) The droplet generator was connected via fluoroethylenepropylene (FEP) tubing (inner diameter 1.0 mm and outer diameter 1.6 mm) to the computer-operated syringe pump, utilizing syringes with volumes of 500 μL (effector solution and cell suspension), 1000 μL (culture medium) and 5000 μL (carrier liquid). Generated segments are transported at a constant flow rate of 200 μL min^-1^ through an optical detector unit for the simultaneous measurement of absorbance and fluorescence data of the droplets. Data were recorded directly through the visually transparent FEP tubing (Fig.1). Absorbance was measured with four light emitting diodes (Agilent, United States) with peak wavelengths of 470, 505, 615 and 750 nm. Measurement of fluorescence was carried out with a laser diode with a peak wavelength of 405 nm (Changchun New Industries Optoelectronics, China) with a longpass filter (425 nm) (Laser Components, Germany) on the detection side. The emitted photons were recorded by a photomultiplier module (Hamamatsu, Japan). To store and incubate the generated droplet sequences, polytetrafluoroethylene (PTFE) tube coils (inner diameter 0.5 mm and outer diameter 1.6 mm) with a length of four meters for 1D- and seven meters for 2D-screening were used.

**Fig. 1.**
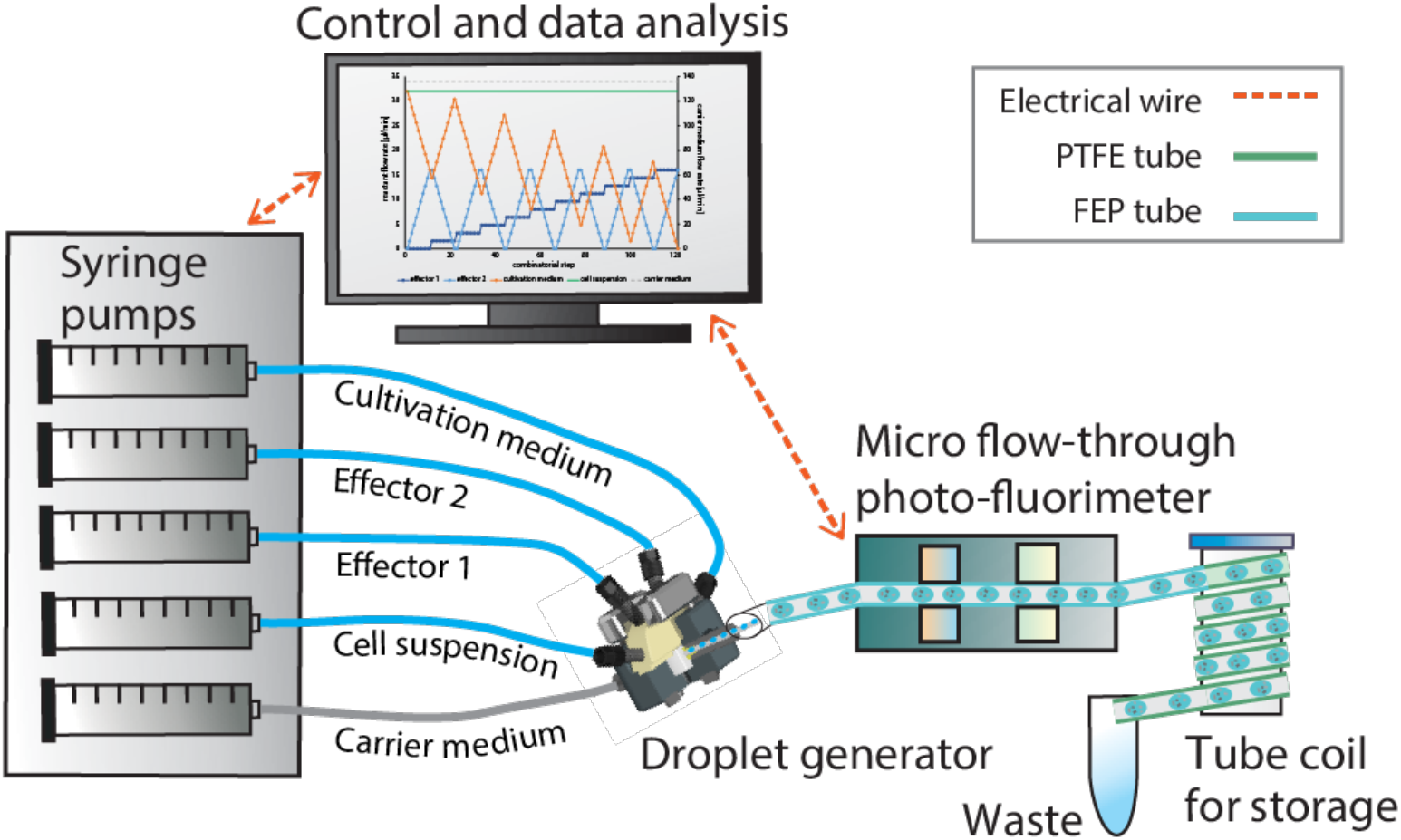
Experimental droplet-based microfluidic setup for 1D and 2D-screening of cultivation parameters used in this study. The illustration on the left shows a 6-channel syringe pump and the droplet generator which generates the droplets. The individual aqueous cell media droplets are separated by the immiscible carrier medium. Droplet size/volume, spacing and composition can be adjusted via the controlled flow rate program of the syringe pump. The generated droplets are measured by a combined photo-fluorimetric micro flow-through detector unit. The droplet sequences are then collected and incubated in PTFE tube coils in an internally illuminated incubator.

#### Experimental Parameters

The syringe pump flow rates of the different fluids were controlled using a LabVIEW program (National Instruments, USA). To adjust the composition of the droplets, the flow rates of the syringe pumps were controlled corresponding to their desired droplet fraction. To investigate the dose-response relationships for single substances (1D-screening), a syringe pump control program with continuous change of the desired effector against a diluting media was used (Fig. 2a). For two effectors (2D-screening), a stepwise increase of the concentrations with a resolution of 10% was applied. Hence, for two effectors, 11 different concentration steps (0, 10, 20, 30, 40, 50, 60, 70, 80, 90, 100%) were combined resulting in 121 different combinations (Fig. 2b). The flow rates of the carrier liquid and the cell suspension were set to 136 μL min–1 and 32 μL min–1 respectively. The flow rates of the effector solutions and cultivation medium were varied within a total flow rate of 32 μL min^-1^. Therefore, the overall flow rate of the segment generation process was kept constant at 200 μL min^-1^. An initial cell density of 5000 cells per 500 nL segment (107 cells mL^-1^) was applied. The generation of the highly resolved 1D- and 2D-screening sequences required approximately four and nine minutes. After droplet formation and initial photo-fluorimetric analysis (t = 0), the droplets were collected in the subsequent connected tube coils for incubation. The tube coils were incubated for 7 days at 30 ± 2°C, 3% CO2 and under 20 ± 5 μmol photons m-2 s-1 illumination. The different droplet sequences were analyzed daily by passing through the photo-fluorimetric detection unit to monitor cell density.

**Fig. 2.**
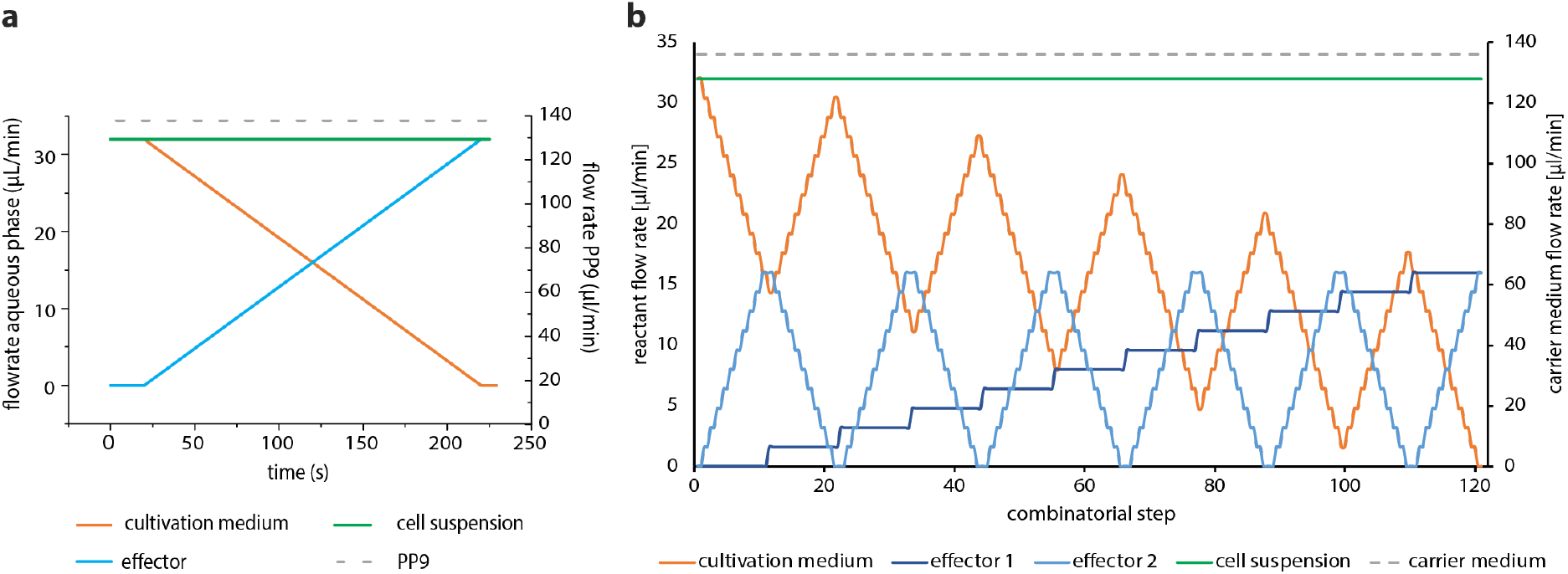
Syringe pump control program for the generation of 1D- and 2D-gradients. **a** Highly resolved dose response screening for single substances (1D-screening). The base cultivation media was continuously substituted over time by the effector-spiked medium and **b** two-dimensional concentration spaces in 11 combinatorial steps, resulting in 121 concentration combinations (2D-screening).

#### Effector screening concentration range

For effector screening, only the concentrations of our test variables were altered, while the remaining medium components were kept at the same concentrations. Stock solutions with two-fold concentrations for 1D screening and four-fold concentrations for 2D screening of the highest final concentration were prepared. These stocks were then used to generate highly resolved concentration gradients using the syringe pump control programs as shown in Fig. 1 and Fig. 2. In all experiments, PP9 (F2 Chemicals Ltd, Lancashire, UK) was used as an immiscible carrier liquid.

#### Data processing

The photo-fluorometric data were recorded with 250 Hz sampling rate. Droplet sequences were measured immediately after formation and after seven days of incubation in the coils (for generation of growth curves, daily measurements were taken). The droplet data were analyzed offline using a custom LabVIEW program which elucidates droplet-specific data from the spectral sensor raw data: number, size, distance between two neighboring droplets as well as extinction and fluorescence measurements. Individual droplets were detected if the absorbance value exceeded the background and achieved a set threshold value. The droplet size correlates with the droplet passage time through the sensor and is determined by the time interval the absorbance exceeds a set threshold value. Due to the inhomogeneous cell distribution in the droplets, growth behavior was analyzed by calculating the integral of the absorbance and fluorescence signal with respect to the droplet size. The recorded autofluorescence intensity of the droplets *I(t)* was normalized to the initial measurement intensity *I(t_0_)* using Eq. 1. Whereas the background was taken on the FEP-tube filled with carrier liquid. Data are given as normalized autofluorescence units (NFU).

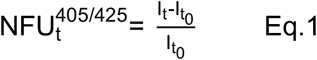

#### Statistics and Reproducibility

In general, for each experiment two droplet sequences were generated and analyzed (Supplementary Fig. 1 red circles and black squares). Each circle represents the intensity for an individual droplet. The droplets were then grouped into 32 concentration ranges from 0 to 100% of effector concentration (corresponding to a 3.1% concentration resolution). The mean and standard deviation were calculated for these 32 concentrations ranges and the resulting curve was plotted as a dose-response curve (Supplementary Fig. 1 blue triangles). For the growth kinetic experiment, each data point represents the average and standard deviation of 50 ± 5 droplets across two independent coils. For 1D- and 2D-screening experiments, a total of 440 ± 21 and 814 ± 26 droplets, respectively, were generated per coil. These droplets were then distributed across the chemical gradients and approximately 10 (1D-screening) and 8 (2D-screening) droplets were analyzed per tested concentration. In sum, each data point represents the average and standard deviation of 20 ± 6 (1D-screening) and 16 ± 6 (2Dscreening) droplets. This redundancy was applied in order to validate the reliability of measurements and to control for stochastic effects that can derive from small reaction volumes and cell numbers. Comparison of final biomass values in the high-density cultivation setup were done with an independent sample Student’s t-test with a significance value of a = 0.05.

## Results and Discussion

In this study we introduce a microfluidics method that allows us to rapidly screen one- and two-dimensional (1D and 2D) culture medium parameters to optimize the growth conditions of industrially relevant cyanobacteria. Our results show that our droplet-based microfluidic approach is well suited to screen culture media conditions in a fast and efficient way, with low material input and a reduced amount of incubator space. In addition, we observed that commonly used medium formulations are not optimized for maximum growth and small changes can result in significantly higher biomass outputs. This method should be widely applicable to a variety of freshwater and saltwater strains and has the potential to facilitate high-throughput strain optimization.

### Microdroplet cultivation setup is suitable to grow freshwater and marine unicellular cyanobacteria

To test the applicability of microdroplet cultivation to cyanobacteria we chose three biotechnologically relevant species. PCC 6803 is a freshwater model cyanobacterium that has been extensively characterized and for which many molecular tools exist. UTEX 2973 is a freshwater fast-growing relative of *Synechococcus elongatus* PCC 7942. Due to their genetic proximity, an average nucleotide identity (ANI) > 99.8%^26^, the tools developed for *Synechococcus elongatus* PCC 7942 are largely transferable to UTEX 2973. UTEX 2973 is of particular interest for biotechnological applications because of its fast growth phenotype with doubling times reported as fast as 1.5 hours^27^. As a third strain, we chose a saltwater strain. A recently isolated strain, *Synechococcus* sp. PCC 11901 (hereafter PCC 11901), is genetically tractable and with clear biotechnological potential due to its reported doubling time of approximately 2 hours and the ability to grow at high light intensities and a range of salinities. Under optimized conditions, PCC 11901 can accumulate up to 33 g dry cell weight per liter. Curiously, it has an ANI of 96.76% when compared to the commonly used *Synechococcus* sp. PCC 7002 strain^20^. Therefore, it may be possible to utilize tools previously developed for *Synechococcus* sp. PCC 7002 in PCC 11901. However, this strain is a vitamin B12 auxotroph. Hence, we decided to use a closely related, spontaneous mutant of this strain, UTEX 3154, that can grow without an external supply of vitamin B_12_.

First, we proceeded to test whether these three cyanobacterial strains could grow in the droplet-based microfluidics setup. For the initial screening, sensors detecting autofluorescence (excitation: 405 nm, emission: 425 nm) and optical density (OD) at 470, 505, 615 and 750 nm were tested over a period of 7 days. In our microfluidic setup, OD reflects the reduction of the intensity of transmitted light by use of a microflow-through photometer. OD typically correlates well with the final cell number. However it does not allow to distinguish between alive and dead cells. Growth can also be monitored by measuring the endogenous cellular autofluorescence with a micro flow-through fluorimeter. The fluorescence can be used to evaluate the approximate number of physiological active cells and, typically, is a more sensitive parameter than OD. The observed increase of the signals in all sensor channels clearly demonstrates cell growth (Supplementary Fig. 2, Supplementary Data Table S1). However, the highest intensities were observed in the OD_470_ (Fig. 3a) and the autofluorescence (Fig. 3b) channels. Therefore, in further experiments, we decided to use the autofluorescence channel to evaluate biomass accumulation. In some cases, biomass accumulation in the droplet storage coils was visible by the naked eye already after 4 days (Fig. 3c). The high cell density was confirmed by the observation of selected individual droplets by light microscopy (Fig. 3d). The typically reported doubling times for PCC 6803 are in the range of 10 to 12 h^28,29^ and approximately 2 h for UTEX 2973^26^ and UTEX 3154^20^. In line with this, our data show that PCC 6803 grows significantly slower than UTEX 2973 and UTEX 3154 (Fig. 3a, 3b). Between UTEX 2973 and UTEX 3154 we observed similar exponential growth rates and final biomass values despite the longer lag phase of UTEX 2973. Overall, the individual growth profiles fit with published literature and show that microdroplet cultivation is suitable for unicellular freshwater and saltwater strains.

**Fig. 3.**
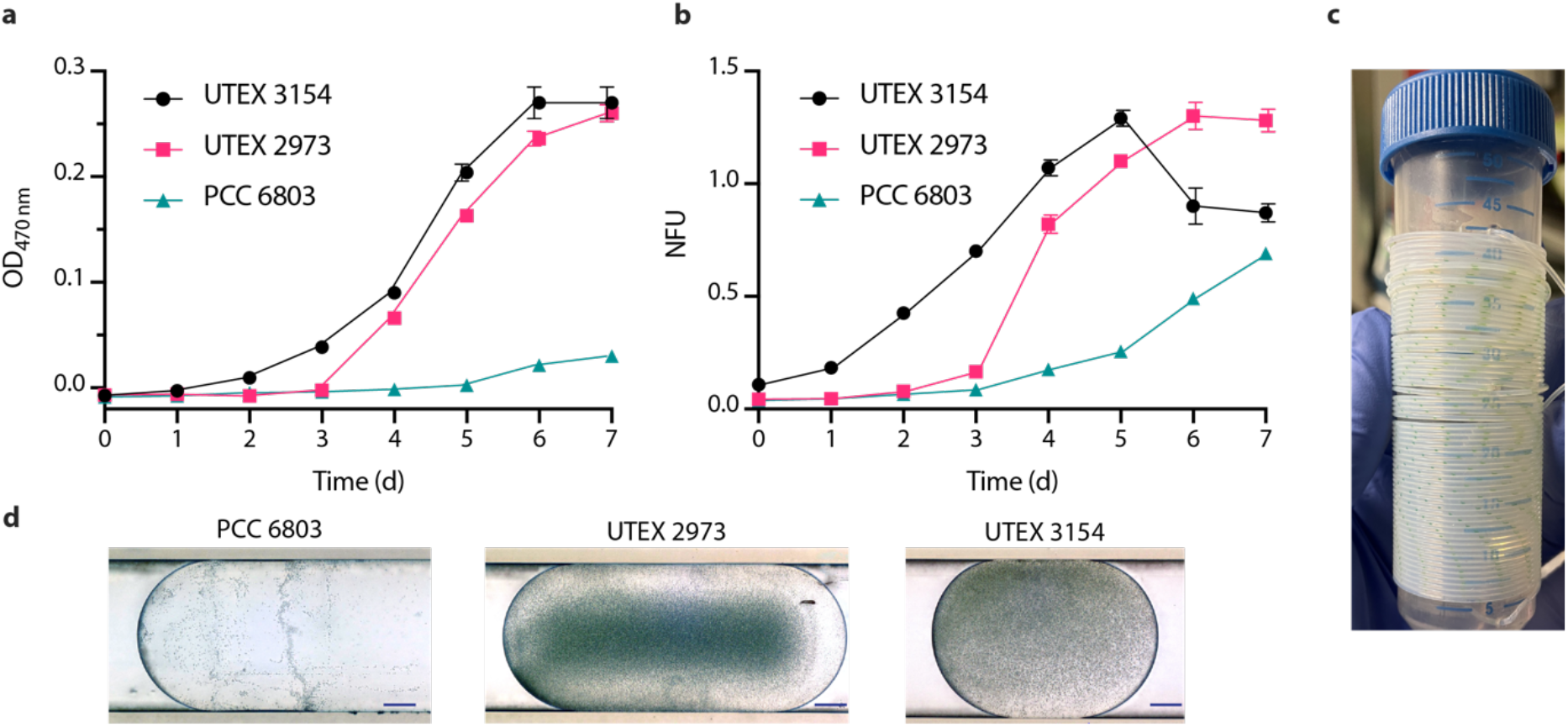
Growth of cyanobacteria in microdroplet setup. Growth kinetics of cyanobacterial strains (UTEX 3154, UTEX 2973 and PCC 6803) in the microfluidic coils measured with multi-channel photo-fluorimeter device showing **a** absorbance (optical density OD) at 470 nm and **b** normalized autofluorescence in the microfluidic setup over a 7 day period. Data points are averages of 50±5 droplets. Error bars represent standard deviation. Normalized autofluorescence is shown as NFU. **c** Image of a microfluidic coil carrying UTEX 2973 droplets. **d** Light microscopy images of cyanobacteria PCC 6803, UTEX 2973 and UTEX 3154 in droplets after 4 days of growth (scale bar: 100 μm).

### Microdroplet technology facilitates high-throughput, high resolution dose response screening

Following the successful droplet-based cultivation of PCC 6803, UTEX 2973 and UTEX 3154, we proceeded to apply the methodology to investigate the response of the cyanobacteria to a variety of medium parameters. Therefore, we designed 1D screening experiments for media optimization. We used the standard growth medium for the respective strains (BG-11 for PCC 6803 and UTEX 2973, AD7 for UTEX 3154) and varied one parameter at a time by microfluidic means. The varied parameters included the nitrogen (N) source, phosphorus (P) source as well as medium salinity and sodium bicarbonate concentrations (Table 1). Furthermore, we proceeded to look at combinatorial effects using 2D screening where both the N and P concentrations were varied simultaneously. An overview with the concentration range of all tested effectors for 1D and 2D-screening experiments is shown in Table 1. A summary of the optimum values for the medium parameters tested in the 1D-screening is shown in Table 2.

**Table 1.** Range of concentrations for 1D- and 2D-screening experiments of selected medium components used in this study. Reference values of concentrations in standard BG-11 TES and AD7 medium are given.

| Medium component | 1D-screening range | 2D-screening range | BG-11 TES medium | AD7 medium |
| --- | --- | --- | --- | --- |
| NaNO<sub>3</sub> (mM) | 2 – 50 | 0 – 30 | 17.6 | 12 |
| K<sub>2</sub>HPO<sub>4</sub> (mM) | 0.05 – 5 | 0 – 0.5 | 0.175 | – |
| KH<sub>2</sub>PO<sub>4</sub> (mM) | 0.05 – 5 | 0 – 0.5 | – | 0.37 |
| NaHCO<sub>3</sub> (mM) | 0 – 100 | – | – | – |
| NaCl (mM) | 0 – 1300 | – | – | 308 |

**Table 2.** Summary of the concentration ranges where the highest growth was observed in the 1D-screening.

| Medium component | PCC 6803 | UTEX 2973 | UTEX 3154 |
| --- | --- | --- | --- |
| NaNO<sub>3</sub> (mM) | 10 | 20 | 30 |
| K<sub>2</sub>HPO<sub>4</sub> (mM) | 2.5 – 3.5 | 0 – 0.5 | – |
| KH<sub>2</sub>PO<sub>4</sub> (mM) | – | – | 0.75 |
| NaHCO<sub>3</sub> (mM) | 50 | NS | NS |
| NaCl (mM) | 100 – 650 | 0 – 300 | 0 - 1100 |

We first started 1D testing of different concentrations of NaNO_3_ ranging from 2 to 50 mM. The data show that the freshwater PCC 6803 and UTEX 2973 achieved maximum biomass values at approximately 10 and 20 mM NaNO_3_, respectively (Fig. 4a, Supplementary Data Table S2). For the saltwater UTEX 3154, the maximum biomass value was achieved at approximately 30 mM NaNO_3_ (Fig. 4a). The typical concentration of NaNO_3_ present in the freshwater cyanobacterial growth medium BG-11 is 17.6 mM. Comparing this value with the limiting NaNO_3_ concentrations for PCC 6803 and UTEX 2973 (10 and 20 mM, respectively), we can conclude that N is not typically the limiting nutrient in BG-11 medium. Accordingly, an earlier study showed that during batch cultivation of PCC 6803 in BG-11 medium one of the major medium limitations may be sulfate ions^30^. Regarding the saltwater medium AD7, the concentration of NaNO_3_ is 12 mM. Considering that the biomass accumulation of UTEX 3154 only peaked at around 30 mM NaNO_3_, our data suggest that N may be a limiting nutrient in AD7. This is supported by the original PCC 11901 strain publication where the authors determined the ideal NaNO_3_ concentration to be between 24 and 48 mM^20^. Overall, these data show that, given enough N, all three cyanobacteria are rapidly limited by other nutrients. Therefore, we proceeded to apply our microfluidic approach to test the effect of varying concentrations of phosphate (K_2_HPO_4_ for UTEX 2973 and PCC 6803 and KH_2_PO_4_ for UTEX 3154) up to a maximum concentration of 5 mM. The data show that the maximum biomass values were achieved between 2.5 and 3.5 mM phosphate for PCC 6803 and UTEX 2973 and 0.75 mM phosphate for UTEX 3154 (Fig. 4b, Supplementary Data Table S3). Looking at the formulations of the base medium, phosphate is present at a concentration of 0.175 mM K_2_HPO_4_ in BG-11 and 0.37 mM KH_2_PO_4_ in AD7. Therefore, our microfluidics growth data suggest that both media are P deficient. This is particularly the case for BG-11 which has a K_2_HPO_4_ concentration 5 to 6 times lower than the levels at which we observed the highest biomass accumulation. It has been suggested that media designed for the growth of algae and cyanobacteria are often P limited due to a lack of compatibility with the Redfield ratio^31^. This ratio describes the amount of carbon (C), N and P typically present in both phytoplankton biomass and in dissolved nutrient pools and has been determined to be 106 C:16 N: 1 P. Based on the media formulation used in the study, AD7 presents a N:P ratio of 32:1 and BG-11 100:1. Thus, supporting our previous hypothesis that both media, and BG-11 in particular, may be P limited.

**Fig. 4.**
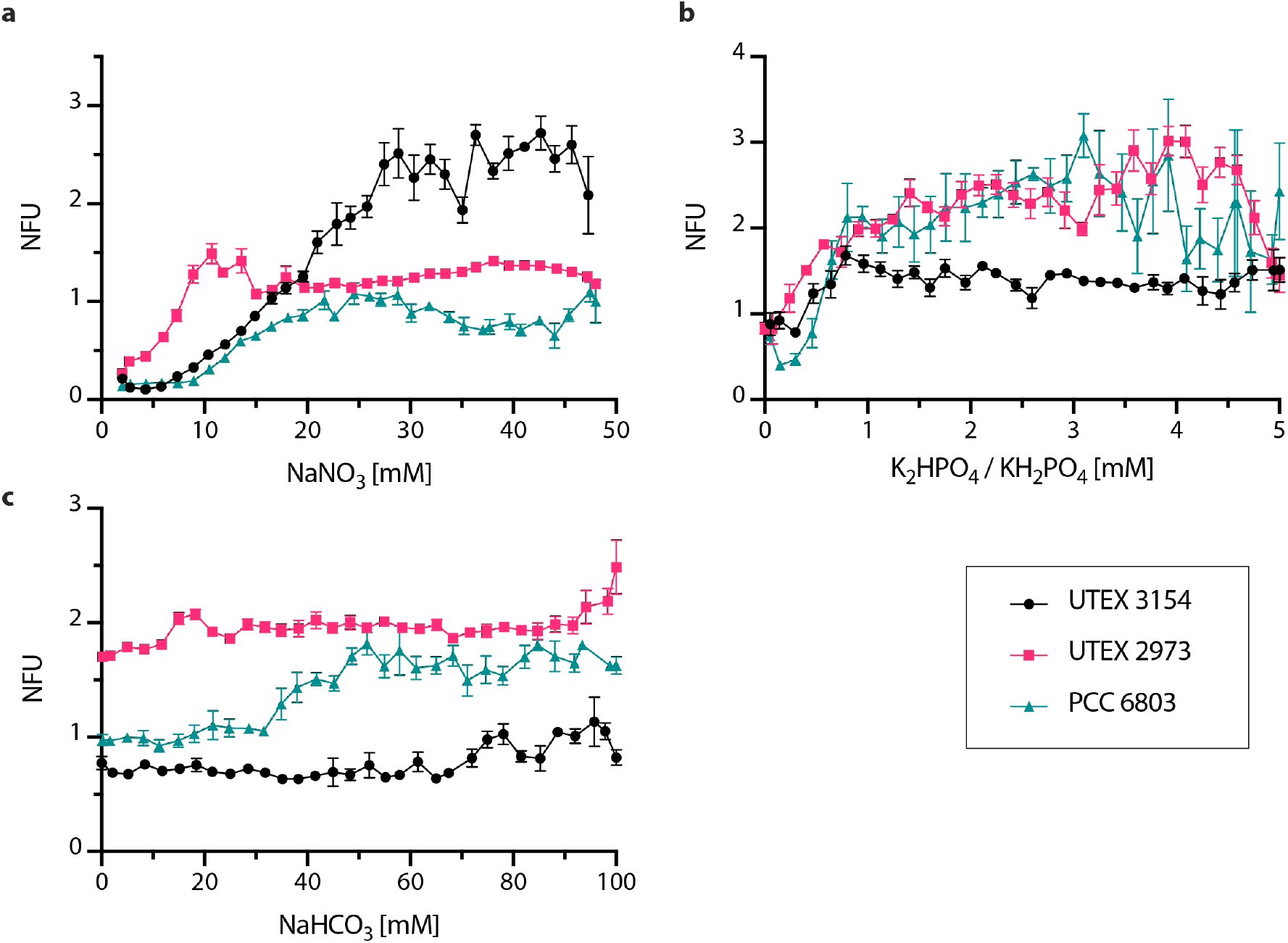
Highly resolved dose-response screenings of key macronutrients in UTEX 3154, UTEX 2973 and PCC 6803. **a** Normalized autofluorescence after 7 days in response to varying concentrations of NaNO_3_. **b** Normalized autofluorescence after 7 days in response to varying concentrations of K_2_HPO_4_ (UTEX 2973, PCC 6803) or KH_2_PO_4_ (UTEX 3154). **c** Normalized autofluorescence after 7 days in response to varying concentrations of bicarbonate (NaHCO_3_). Data points are averages of 10±3 droplets. Error bars represent standard deviation. Normalized autofluorescence is shown as NFU.

The autotrophic growth of cyanobacteria requires a source of inorganic C. This is typically provided as CO_2_ which can be supplied by air (0.04% CO_2_), CO_2_ enriched air (typically 1-5%) or in the form of bicarbonate salts. Installing a gas supply to cyanobacterial cultures can be costly and presents a logistical challenge for parallel experimentation. Therefore, sodium bicarbonate is a popular low-cost inorganic C source for cyanobacterial medium. Here, we tested the addition of NaHCO_3_ to BG-11 and AD7 in the range of 0 to 100 mM (Fig. 4c, Supplementary Data Table S4). The addition of bicarbonate improved the growth of PCC 6803 from 35 mM and achieved a maximum biomass accumulation at approximately 50 mM. For UTEX 2973 and UTEX 3154 there was no significant difference in biomass accumulation in the tested range. In our previous experiments we have shown that BG-11 is P limited (Fig. 4b). This could explain the lack of response of UTEX 2973 to the varying bicarbonate concentrations. In regard to UTEX 3154, a previous study in *Synechococcus* sp. PCC 7002 showed that significant biomass accumulation was only visible with bicarbonate concentrations higher than 500 mM^32^. This suggests that the range tested here may not comprise the ideal bicarbonate values for this cyanobacterium. However, *Synechococcus* sp. PCC 7002 and UTEX 3154 present significant genetic differences therefore it is difficult to make definite conclusions. Finally, another factor could be light limitation. In our experiments we used 20±5 μmol photons m^-2^ s^-1^ while the maximum growth rates of UTEX 2973 were reported at 1500 μmol photons m^-2^ s^-1^, 42°C and 5% CO_2_^27^. However, this is not directly comparable as the light path in our microfluidic setup is 1.0 mm as opposed to the 27 mm used to determine the maximum growth rates of UTEX 2973 in the literature. Further tests would be needed to establish a robust comparison.

Salinity of the growth medium is a critical factor as high salinities can induce a variety of stresses and consequently pose a challenge to cell survival^33^. In addition, future large-scale cultivation of cyanobacteria should be done in seawater due to the limited freshwater resources present on Earth. Therefore, there has been an increased interest in prospecting for and developing salt tolerant cyanobacterial chassis. Here, we used our microfluidics platform to determine the salt tolerance of PCC 6803, UTEX 2973 and UTEX 3154 (Fig. 5, Supplementary Data Table S5). As expected, the two freshwater strains exhibit lower salt tolerances than UTEX 3154. UTEX 2973 exhibits the lowest salt tolerance with a decline in biomass accumulation starting at 0.3 M NaCl. PCC 6803 maintains similar levels of biomass accumulation until 0.65 M with a sharp decline observed thereafter. Total inhibition was observed of both PCC 6803 and UTEX 2973 at 0.7 M NaCl. UTEX 3154 accumulates similar biomass levels until approximately 0.8 M, whereafter a sharp decline is also observed. It is worth noting here that the base AD7 used in this study contains 308 mM NaCl and 8 mM KCl therefore the data show that UTEX 3154 can tolerate salt concentrations up to 1.1 M NaCl. These values are in accordance with the published literature on salt tolerance in cyanobacteria^34^. Our data show that this microfluidics setups can be used for high-resolution screening for optimum salinity cultivation conditions of single-celled cyanobacteria and could serve as an effective high-throughput method to screen for strains with increased salt tolerance in future studies.

**Fig. 5.**
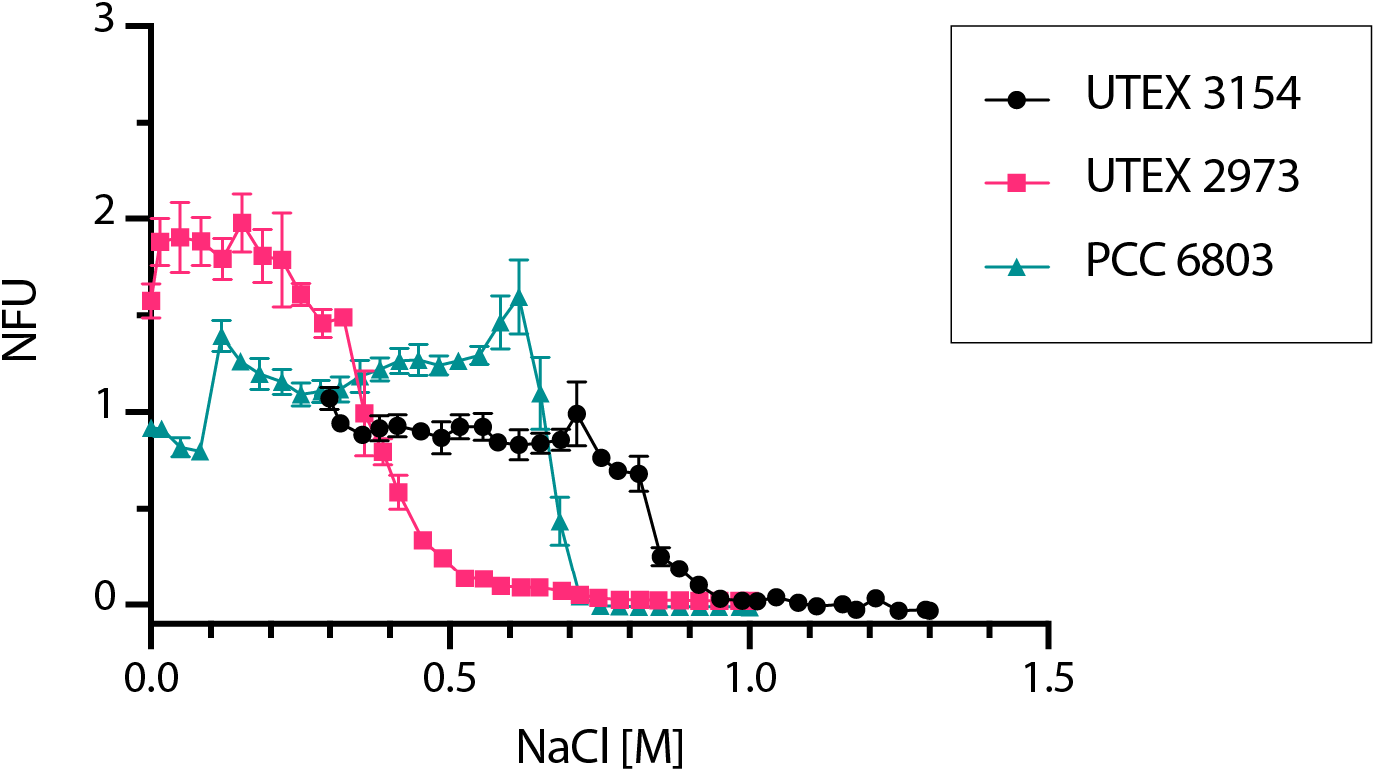
1D screening of NaCl tolerance in cyanobacteria. Normalized autofluorescence after 7 days in response to varying concentrations of NaCl shown for three different cyanobacterial strains, UTEX 3154, UTEX 2973 and PCC 6803. Data points are averages of 10±3 droplets. Error bars represent standard deviation. Normalized autofluorescence is shown as NFU.

### Combinatorial effect of phosphorus and nitrogen on the growth behavior of cyanobacteria

The above results demonstrate that our microfluidics platform can rapidly generate droplet screening sequences with a wide range of varying parameters (i.e. 1D screening). However, it is often of interest to screen a combination of parameters. Therefore, we took advantage of the capabilities of the droplet-based technique to simultaneously vary two independent parameters (2D screening). The result of a 2D media optimization experiment are 2D response diagrams where we can easily estimate the effect of the two variables on one readout parameter (e.g. biomass accumulation). For the 2D proof of concept experiment we decided to screen N (NaNO_3_) and P (K_2_HPO_4_ or KH_2_PO_4_) sources in conjunction (Fig. 6, Supplementary Data Table S6, S7, S8). These two parameters were chosen because the N:P ratio is one of the key parameters that influences algal growth and rapidly optimizing N and P concentrations is fundamental for cost-efficient scale-up of microalgal cultures^35^. Our data show that final biomass values in PCC 6803 and UTEX 2973 peaked at the maximum N values tested (0.45 mM K_2_HPO_4_/30 mM NaNO_3_ for PCC 6803 and 0.4 mM K_2_HPO_4_/30 mM NaNO_3_ for UTEX 2973) (Fig. 6a, b). Interestingly, an increase in N at low P values or an increase of P at low N values was not sufficient to obtain high biomass values. It was only when both parameters were increased simultaneously that a significant increase in final biomass values was observed. This contrasts with the conclusions from the 1D screening data where P seemed to be limiting the culture. This suggests that it is more efficient to find a good balance between N and P instead of just increasing one parameter. Therefore, 2D screening is fundamental to understanding nutrient dynamics in cyanobacterial cultures. Regarding the N:P ratio, both strains achieved their maximum biomass values at a ratio of approximately 100:1. This is the same ratio as BG-11 which suggests that the classic cyanobacterial medium has a good N:P ratio but would benefit from higher concentrations of both N and P. For UTEX 3154 no clear trend was observed within the tested N:P range (Fig. 6c). Together with the 1D screening data, this suggests that N may be the limiting nutrient in the AD7. However, the possibility that a nutrient other than N or P is limiting the culture remains open.

**Fig. 6.**
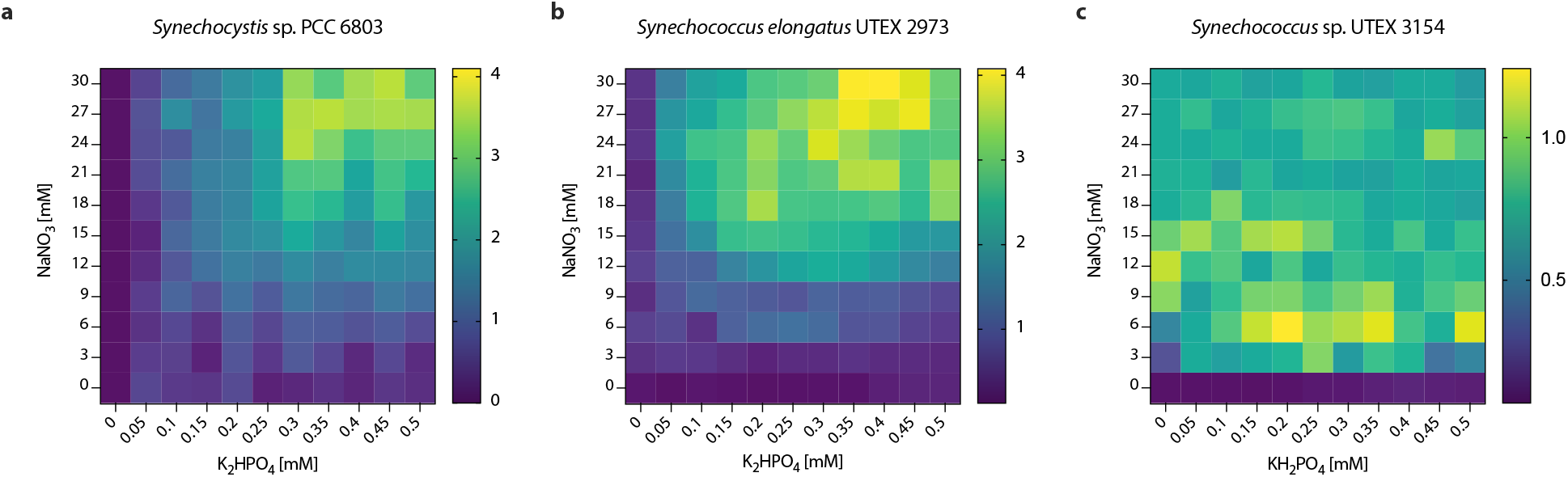
2D-screening of nitrogen and phosphorus source in cyanobacterial cultivation using standard growth media. **a** PCC6803 normalized autofluorescence after 7 days in response to varying concentrations of NaNO_3_ and K_2_HPO_4_ in BG-11 medium. **b** UTEX 2973 normalized autofluorescence after 7 days in response to varying concentrations of NaNO_3_ and K_2_HPO_4_ in BG-11 medium. **c** UTEX 3154 normalized autofluorescence after 7 days in response to varying concentrations of NaNO_3_ and KH_2_PO_4_ in AD7 medium. Data points are averages of approximately 5±3 droplets.

### Improved N:P ratios from 2D screening increase biomass accumulation in high density cultivation

To assess whether the optimal N:P concentrations suggested by the microfluidic 2D experiments can be translated to a larger scale cultivation setup, we carried out growth assays in a laboratory setting. To this end, PCC 6803 and UTEX 2973 were grown in 25 mL high-density cultivators where gaseous CO_2_ is supplied via integrated semi-permeable membranes. With this setup, growth was compared in standard BG-11 medium (0.175 mM K_2_HPO_4_, 17.6 mM NaNO_3_) and BG-11 with the optimized N:P concentrations obtained from the 2D N:P microfluidic screening experiment. More specifically, 0.45 mM K_2_HPO_4_ with 30 mM NaNO_3_ for PCC 6803 and 0.4 mM K_2_HPO_4_ with 30 mM NaNO_3_ for UTEX 2973. For both PCC 6803 and UTEX 2973 significant increases (p < 0.05) of final biomass values of 7.4% (Fig. 7a) and 15.7% (Fig. 7b), respectively, could be observed. Overall, this confirmed that the findings from the microfluidic experiments are transferable to biotechnologically relevant high-density cultivation setups.

**Fig. 7.**
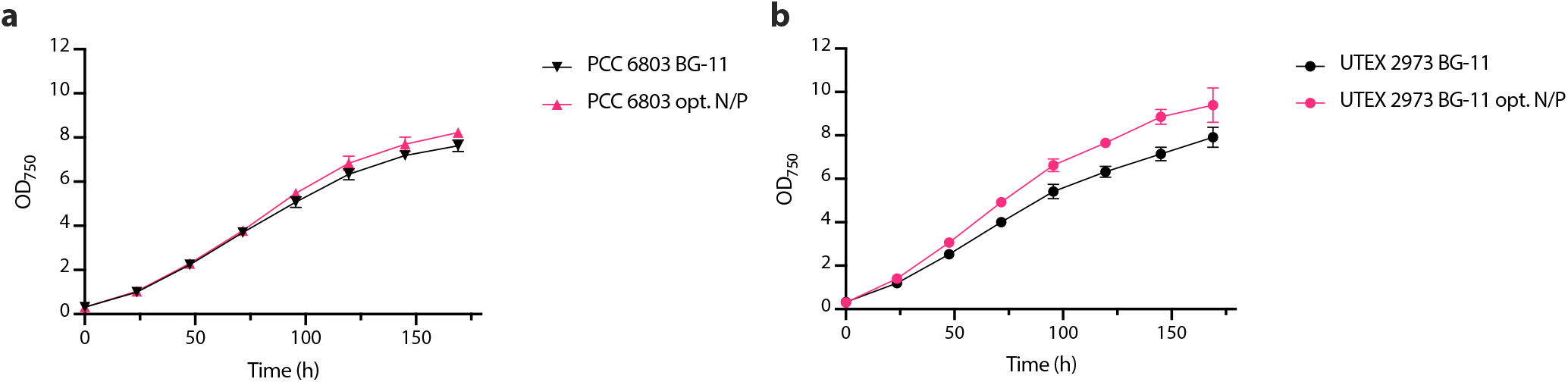
High-density cultivation in BG-11 medium containing N:P ratios optimized through 2D microfluidic screening. Growth of **a** PCC 6803 and **b** UTEX 2973 in standard vs. optimized N/P ratio observed by measuring optical density (OD) at 750 nm. n=3 and error bars represent standard deviation.

## Conclusions

To achieve cost-effective scale-up of cyanobacteria it is crucial to screen for the ideal cultivation conditions. In particular, medium optimization can significantly increase biomass and product yields. However, the cost and time investment required to test different levels of key nutrients, and the possible interactions amongst them, is prohibitive with standard reactor setups. Recently, miniaturized multiplex screening has been suggested as a route to tackle the large parameter fields of media optimization. In this study we used a droplet-based microfluidic technique to improve the cultivation of three different biotechnologically relevant cyanobacterial strains. Our proof of concept demonstrated that the strains could be successfully cultivated, and growth data could be collected online with non-invasive, in-situ measurements. The 1D-screening data confirmed that our microfluidic platform is well-suited for the investigation of cyanobacterial response towards single nutrients. In addition, the 2Dscreening allowed us to easily explore the two-dimensional space of nutrient interactions. In conclusion, this study shows that microfluidics can play a valuable role in improving the costeffectiveness of cyanobacterial cultivation. We also expect that this microfluidics approach can be generalized to other applications such as expression level optimization of engineered cyanobacteria and bioprospecting.

## Author Contributions

Conceptualization and study design: JC, DAR and JAZZ. Experimental work and data analysis: JC, DAR, TX and JAZZ. Microfluidic device and system development: GAG, JC. Writing – original draft: JC, DAR and JAZZ. Writing – review & editing: JC, DAR, GAG and JAZZ. Funding acquisition and project administration: JC, DAR, GAG and JAZZ. All authors have read and approved the final manuscript.

## Acknowledgments

We thank Prof. Michael Köhler for the fruitful discussion. We thank Frances Möller and Steffen Schneider for lab assistance. JC gratefully acknowledges financial support from the project “Screen | in drop lines” (2016FE9016) by Thüringer Aufbaubank and a habilitation scholarship from the TU Ilmenau, DAR was supported by the Alexander von Humboldt Foundation and by the IMPULSE^*project*^ (IP-2020-03, Friedrich Schiller University Jena). This work was funded by the Deutsche Forschungsgemeinschaft (DFG, German Research Foundation) under Germany’s Excellence Strategy – EXC 2051 – Project-ID 390713860 (JAZZ) and the by the Deutsche Forschungsgemeinschaft (DFG, German Research Foundation) – CRC 1127/2 – 239748522 (JAZZ).

## Data availability

All cyanobacterial strains used in this study are available for purchase from the Pasteur Culture Collection, Paris (France) or the UTEX Culture Collection of Algae, University of Texas, Austin (USA). The raw data of all experiments are provided in the Supplementary Data files.

