## Supplementary Material for "Droplet-based microfluidics as a media optimization tool for cyanobacteria"

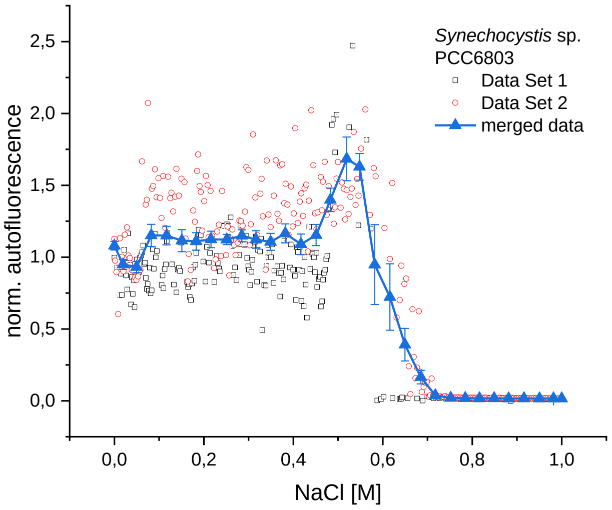


**Supplementary Fig. 1** A representative example of how the microdroplet raw data is visualized and treated for each experiment. Details can be found in the Materials &Methods section “Statistics and Reproducibility”.

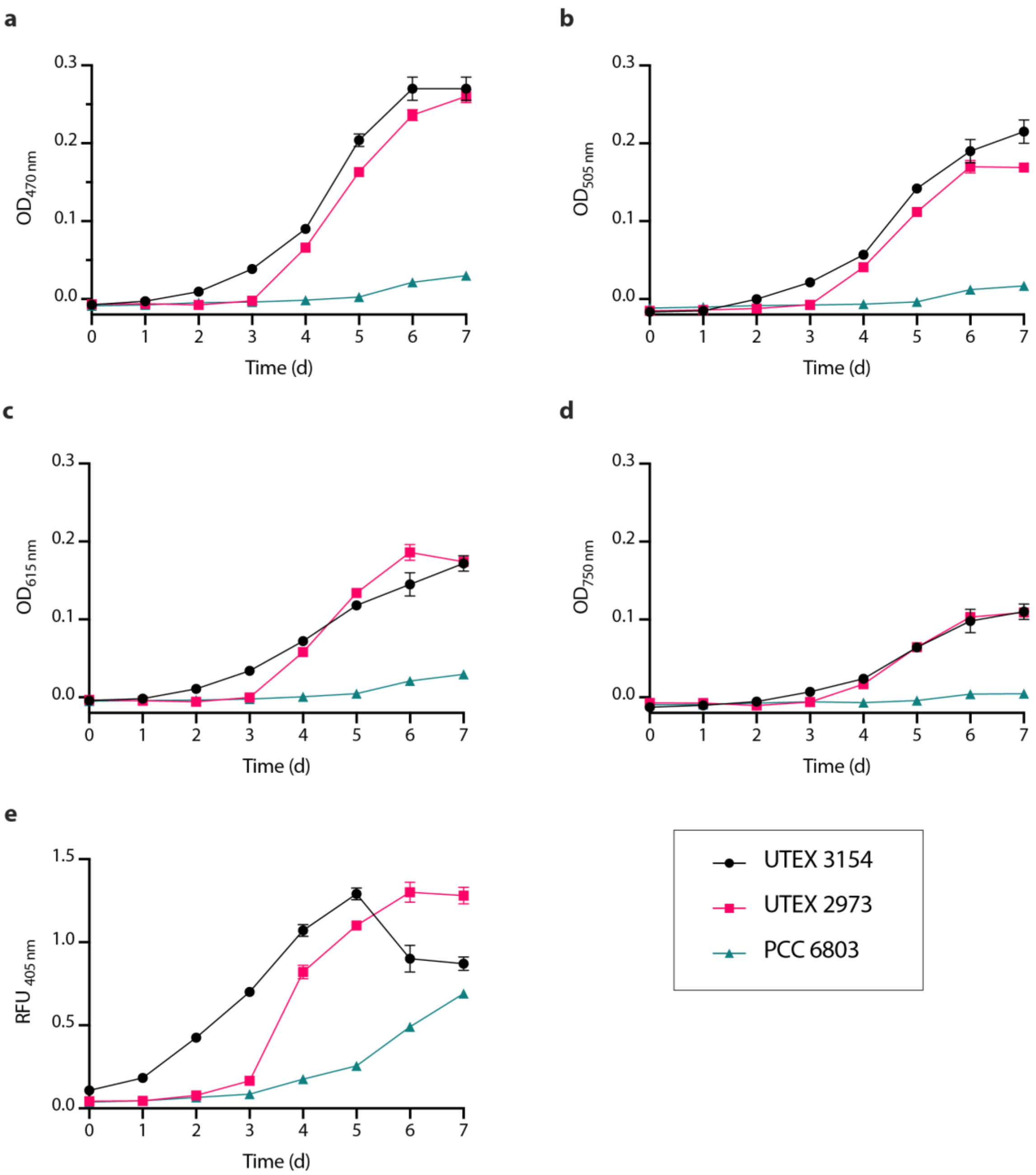


**Supplementary Fig. 2** Growth kinetics of three cyanobacterial strains in microdroplet setup over a period of 7 days. Measurements of multi-channel photofluorimeter with different exctinction wavelengths and fluorescence. **a** optical density (OD) at 470 nm, **b** OD at 505 nm, **c** OD at 615 nm, **d** OD at 750 nm and **e** fluorescence (excitation: 405 nm, emission: 425 nm). Data points are averages of approximately 50 droplets. Error bars represent standard deviation.
